# Inactivation of NMDAR and CaMKII signaling within the prelimbic cortex blocks incubated cocaine- and sucrose-craving

**DOI:** 10.1101/2025.07.10.664193

**Authors:** Laura L. Huerta Sanchez, Natasha M. Siao, Sanil R. Chaudhari, Julie E. Barrios, Audrey Y. Na, Mirette G. Tadros, Megan L. McConnell, Rachel M. Kaplan, Caden R. Lane, Hoa H.T. Doan, Ashley B. Liger, Tessa C. Chou, Serena Marcon, Fernando J. Cano, Tod E. Kippin, Karen K. Szumlinski

## Abstract

The incubation of craving is a term coined to characterize the behavioral phenomenon wherein cue-elicited craving strengthens over a period of abstinence. Incubated cocaine-craving is mediated, at least in part, by increased glutamate release within the prelimbic cortex (PL). We hypothesized that this glutamate release stimulates NMDA-type glutamate receptors (NMDARs) leading to calcium-dependent activation of CaMKII signaling that drives incubated craving. To test this hypothesis, adult male and female Sprague-Dawley rats were trained to self-administer either IV cocaine or sucrose pellets (6h/day x10 days) and tested for cue-elicited cocaine- or sucrose-craving in early versus later (i.e. after incubation) withdrawal. Incubated cocaine-seeking was associated with increased CaMKII activity in the PL, but no change in NMDAR subunits. In contrast, incubated sucrose-craving was associated with many sex-dependent changes in both NMDAR subunit expression and CaMKII activation that were subregion-selective. An intra-PL infusion of the NMDA antagonist D-AP5 (2.5 or 7.5 µg/side) or the CaMKII inhibitor myr-AIP (10 pg/side) blocked both incubated cocaine- and sucrose-craving, with no effects detected in early withdrawal. Co-infusion of both D-AP5 and myr-AIP exerted an additive effect on incubated cocaine-craving that was larger than either antagonist alone. These data corroborate earlier evidence for distinct biochemical correlates within mPFC between incubated cocaine- and sucrose-craving and, for the first time, demonstrate that NMDARs and CaMKII activation within the PL are common drivers of incubated craving that operate via independent signaling pathways suggesting combined pharmacological treatments may have greater efficacy in managing addiction.

## INTRODUCTION

Cocaine Use Disorder (CUD) is a chronic relapsing disorder responsible for an estimated 27% of the 107,543 overdose deaths in 2023 [1]. One major trigger for relapse in CUD is drug-associated cues and cue-reactivity strengthens during periods of abstinence [2,3]. This “incubation of craving” phenomenon [4] is observed in both humans diagnosed with, and in animal models of, CUD [c.f., 5]. Cue-elicited craving is associated with changes in prefrontal cortex (PFC) activity in humans [6–8] and neural measures of cocaine cue-reactivity incubate in individuals with CUD [3]. Consistent with this, animal models of incubated craving show increased indices of cellular activity within the ventromedial aspect of the PFC (vmPFC)--which the prelimbic (PL) and infralimbic (IL) cortices) [9–11]--that we theorize is driven by cue-elicited glutamate release within the PL in late withdrawal [12,13].

The receptors mediating the glutamate-dependent drive on incubated responding are presently not clear. However, AMPA/kainate receptor (AMPAR/KAR) activation within the PL is required for the expression of incubated cocaine-craving in rats [14]. In addition, N-methyl-D-aspartate receptors (NMDARs) also mediate “fast” excitatory synaptic transmission that could incubate craving. NMDARs are comprised of 7 different subunits; GluN1 is the obligatory NMDA receptor subunit, required for channel function, while GluN2 subunits influence receptor scaffolding, agonist sensitivity, pharmacology, conductance and desensitization [15–17]. NMDAR stimulation activates Ca^2+^-dependent intracellular signaling. While we know that (1) both GluN2A and GluN2B subunit expression are increased in dorsomedial aspects of the PFC (dmPFC) of cocaine-experienced rats during late withdrawal [18,19] and (2) GluN3-containing NMDARs within the nucleus accumbens (NAc) are required for the expression of incubated cocaine-craving in rats [20], the functional relevance of NMDAR activation within vmPFC subregions for incubated cocaine-craving has not been explored to the best of our knowledge.

A major downstream effector of NMDAR activation is calcium/calmodulin-dependent protein kinase II_*α*_ (CaMKII) [21]. CaMKII plays a key role in synaptic plasticity, learning, and memory [22–24], notably within the context of substance use disorders [25–28], including incubated cocaine-craving [29]. Increased expression of p(Thr286)-CaMKII within the PL is observed in male rats exhibiting incubated cocaine-craving following short-access cocaine self-administration procedures [30], implicating autonomous CaMKII activation [31] within the PL in incubated cocaine-craving. Whether increased CaMKII activation (1) extends to rats with more extensive cocaine-taking experience, to female rats and to non-drug reinforcers (e.g. sucrose), (2) relates to changes in NMDAR subunit expression or (3) is functionally relevant for incubated craving are unknown. Thus, we examined the role for NMDA-CaMKII signaling within mPFC subregions in the expression of incubated cocaine- and sucrose-craving in male and female rats. Our results confirm elevated p(Thr286)-CaMKII expression in the PL as a biochemical correlate of incubated cocaine-seeking, independent of changes in total NMDAR subunit expression. Incubated sucrose-seeking was associated with sex- and subregion-dependent changes in NMDAR subunit expression and CaMKII activation that were distinct from cocaine. Irrespective of sex, pharmacological inhibition of NMDARs or CaMKII reduced the magnitude of incubated cocaine- and sucrose-craving. The effect of inhibitor co-infusion on incubated cocaine-craving was larger than that of either inhibitor alone, indicating that CaMKII activation by NMDARs and other upstream mediators drive this phenomenon. The present results further support that the biochemical correlates of incubated cocaine- and sucrose-seeking are distinct with respect to direction of effect, as well as their sex- and subregional-specificity [14,30,32]. Despite this, both NMDAR and CaMKII signaling within the vmPFC is functionally relevant for incubated craving for both a drug and non-drug reinforcer of relevance to our neurobiological understanding of incubated craving.

## METHODS

Detailed methods are in the Supplemental Material.

### Subjects, Surgery and Self-Administration Procedures

Adult male and female Sprague Dawley rats (225-250 g; Charles River Laboratories, Hollister, CA, USA), were housed under a reverse light-dark cycle (lights off:10:00 am), with *ad libitum* access to food and water. Rats were implanted with implant bilateral guide cannulae over the PL and an intravenous (IV) catheter into the jugular vein (for IV cocaine reinforcement). Rats received either IV cocaine or 45 mg banana-flavored sucrose pellet reinforcement (10 days, 6 hours per day) paired with a light+tone CS.

### Tests for Cue-elicited Craving

As conducted previously [10,11,14,30,32–35], rats employed for immunoblotting underwent a 2-h test for cue-elicited craving (Cue Test) in either early withdrawal (WD1 for sucrose; WD3 for cocaine) or on withdrawal day 30 (WD30), while rats employed in the neuropharmacological studies underwent a 30-minute Cue Test on either WD1 or following at least 30 days of withdrawal (WD30+). In all cases, pressing on the active, formerly reinforced, lever resulted in the presentation of the tone-light conditioned stimulaus, but no reinforcer delivery. Upon the completion of the 2-h session, vaginal smears were collected (females only) to permit estrous cycle phase determination [14] and the PL and IL subregions were dissected over ice and stored at −80 ^0^C until processed.

To investigate the role of NMDA and CaMKII signaling in incubated craving, rats received bilateral PL infusions (0.5 µl/minute rate for 1 minute) of either 2.5 or 7.5µg/side of D-2-Amino-5-phosphonopentanoic acid (D-AP5; Tocris, Minneapolis, MN) [36,37], 10 pg/side of the CaMKII inhibitor myristoylated Autocamtide-2-related inhibitory peptide (myr-AIP; Tocris, Minneapolis, MN) [38,39], or vehicle (VEH). To investigate NMDA-CaMKII signaling pathway interactions, rats were co-infused with 7.5µg/0.25µl/side D-AP5 and 10 pg/0.25 µl/side myr-AIP (total volume=0.5 µl/side). Immediately following microinjection, rats underwent a 30-min Cue Test and a second identical Cue Test the next day to examine for any carry-over effects of our neuropharmacological manipulations [11,32,35]. Brains were extracted and drop-fixed 4% paraformaldehyde, prior to sectioning (30 μm thick) and histological verification of cannula placement using Nissl staining procedures.

### Immunoblotting

Immunoblotting was conducted on the PL and IL of rats exhibiting incubated cocaine- and sucrose-craving. Procedures for preparing tissue homogenates, detecting, and quantifying protein expression are similar to those conducted in our laboratory previously [14,30–32].

## RESULTS

### Operant-conditioning for cocaine and sucrose reinforcement

The results related to the operant-conditioning phase of each study are summarized in **Table 1**.

**Table 1:** Means ± SEMs of the number of active and inactive lever-presses, as well as reinforcers earned, by male and female rats over the last 3 days of the cocaine or sucrose self-administration phases of the different experiments in this report.

**Figure Legends**
|  | ACTIVE | REINFORCER | INACTIVE |
| --- | --- | --- | --- |
| **Exp. 1—Immunoblotting: Incubated Cocaine-Craving** |  |  |  |
| **Male:** |  |  |  |
| WD3 Control | 26.55±8.72 | 18.83±1.99 | 12.87±5.54 |
| WD3 Cocaine | 64.06±11.88 | 52.53±2.42 | 6.23±9.48 |
| WD30 Control | 18.15±5.21 | 14.92±1.65 | 10.35±3.90 |
| WD30 Cocaine | 81.4±12.47 | 74.47±2.35 | 7.48±12.37 |
| **Female:** |  |  |  |
| WD3 Control | 28.17±11.07 | 7.9±1.91 | 5.47±1.14 |
| WD3 Cocaine | 70.4±9.69 | 64.93±0.73 | 3.73±9.69 |
| WD30 Control | 7.73±1.30 | 13.85±0.59 | 2.73±6.96 |
| WD30 Cocaine | 70.74±7.67 | 65.15±11.18 | 18.15±5.71 |
| **Exp. 2—Effect of intra-PL AP5 on incubated cocaine-craving** |  |  |  |
| WD1 VEh | 116.13±25.31 | 96.60±21.87 | 9.96±2.32 |
| WD30 VEh | 102.2±10.66 | 80.20±6.97 | 8.66±8.83 |
| WD30 AP5 (2.5 μg) | 87.74±14.81 | 70.86±11.01 | 6.40±9.05 |
| WD30 AP5 (7.5 μg) | 98.15±13.37 | 85.37±10.12 | 4.03±16.61 |
| **Exp. 3—Effect of intra-PL AP5 on incubated sucrose-craving** |  |  |  |
| WD1 VEh | 237.21±15.71 | 177.5±5.54 | 20.8±4.10 |
| WD30 VEh | 199.83±13.52 | 178.46±9.50 | 12.83±2.07 |
| WD30 AP5 (7.5 μg) | 198.71±11.62 | 187.56±8.26 | 6.93±1.32 |
| **Exp. 4—Effect of intra-PL myr-AIP on incubated cocaine-craving** |  |  |  |
| WD1 VEh | 203.08±68.93 | 133.42±51.09 | 13.17±4.50 |
| WD30 VEh | 205.63±105.89 | 161.17±10.10 | 14.00±14.13 |
| WD30+ myr-AIP and AP5+myr-AIP | 240.06±53.98 | 111.17±24.32 | 10.17±1.32 |
| **Exp. 5—Effect of intra-PL myr-AIP on incubated sucrose-craving** |  |  |  |
| WD1 VEh | 212.56±9.84 | 162.06±9.10 | 15.83±6.31 |
| WD40 VEh | 212.58±9.59 | 173.38±16.11 | 5.42±1.36 |
| WD40 myr-AIP | 213.04±19.56 | 175.54±6.43 | 6.95±2.62 |
| **Exp. 6—Effect of intra-PL AP5 and myr-AIP in early cocaine withdrawal** |  |  |  |
| WD1 VEh | 115.16±9.84 | 67.47±21.69 | 11.80±2.85 |
| WD1 AP5 (μg) | 106.30±17.51 | 97.88±11.42 | 9.58±3.42 |
| WD1 myr-AIP | 113.24±16.61 | 99.50±14.39 | 7.47±2.70 |
| **Exp. 7—Immunoblotting: Incubated Sucrose-Craving** |  |  |  |
| **Male:** |  |  |  |
| WD1 | 197.91±11.40 | 170.64±11.54 | 12.33±2.68 |
| WD30 | 170.57±10.08 | 140.51±7.59 | 15.55±3.02 |
| **Female:** |  |  |  |
| WD1 | 184.4±12.00 | 160.22±9.04 | 9.95±3.31 |
| WD30 | 176.55±10.43 | 140.52±9.26 | 16.64±4.52 |

### Protein correlates of incubated cocaine-craving

The results for the tests for cue-elicited cocaine-craving are summarized **Fig.1A** and detailed in [14], with the PL protein expression results provided below. The IL results, as well as the relationship of PL and IL results to estrous cycle phase, are both provided in the Supplementary Materials and depicted in **Suppl. Fig.2-3**.

**Figure 1:**
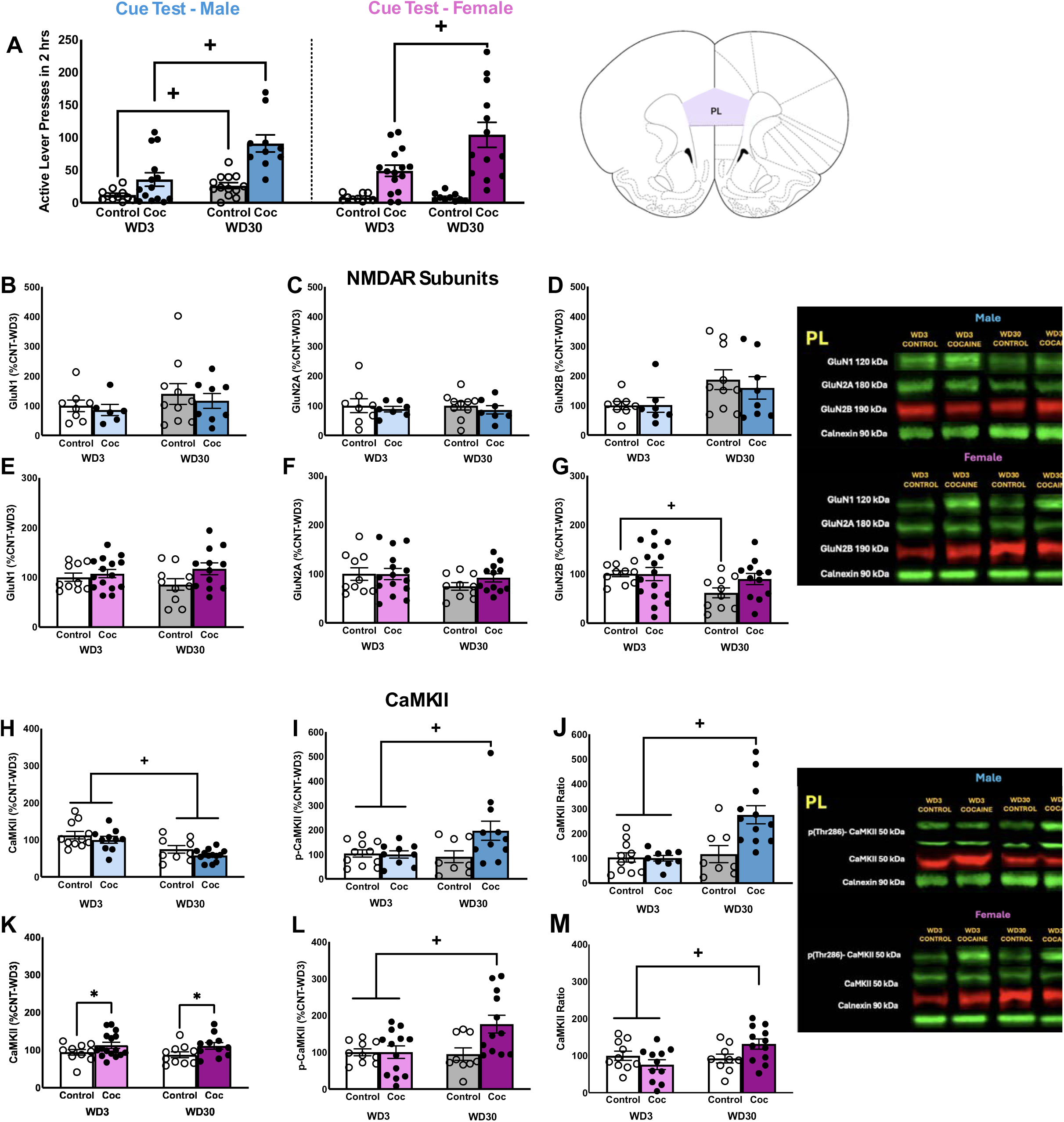
Immunoblotting conducted on the PL subregion of cocaine-incubated rats. **(A)** Active lever-pressing exhibited by male and female rats tested for incubated cocaine-craving. Details of the statistical analyses are provided in Ref. 14. Immunoblotting for the total protein expression of **(B)** GluN1 [F(1,44)<0.026, p>0.873], **(C)** GluN2A [F(1,44)<0.020, p>0.888] and **(D)** GluN2B [F(1,44)<0.002, p>0.968] within the PL of male rats indicated no differences between cocaine-experienced rats (Coc) and cocaine-naive controls (Control), when examined on withdrawal days 3 or 30 (respectively, WD3 and WD30). Comparable immunoblotting conducted on the PL of females also indicated no group differences in **(E)** GluN1 [F(1,47)<3.731, p>0.060] or **(F)** GluN2A [F(1,47)<2.424, p>0.127], while a Withdrawal effect was detected for GluN2B **(G)** [F(1,47)=1.380, p=0.05]. Immunoblotting for the total protein expression of **(H)** CaMKII, **(I)** p(Thr286)-CaMKII and **(J)** their relative expression within the PL subregion of male rats. In contrast to Controls, cocaine-experienced males exhibited a time-dependent increase in both total [for Controls: t(1,17)=0.503,p=0.622; for Cocaine: t(1,18)=2.119, p=0.048] and relative expression of p(Thr286)-CaMKII [for Control: t(1,17)=-0.373, p=0.714; for Cocaine: t(1,19)=-4.017, p<0.001]. Comparable immunoblotting conducted on PL tissue from females **(K-M)** revealed a similar pattern of group differences for both total [for Controls: t(1,17)=0.265, p=0.794; for Cocaine: t(1,23)=-2.623, p=0.015] and relative expression of p(Thr286)-CaMKII [for Control: t(1,17)=0.454, p=0.656; for Cocaine: t(1,22)=2.508, p=0.020]. Representative immunoblots are provided for all proteins examined. The data represent the means ± SEMs of the individual animals indicated. *p<0.05 vs. Control (main Drug effect); +p<0.05 vs. WD3 (Withdrawal effect).

**Figure 2:**
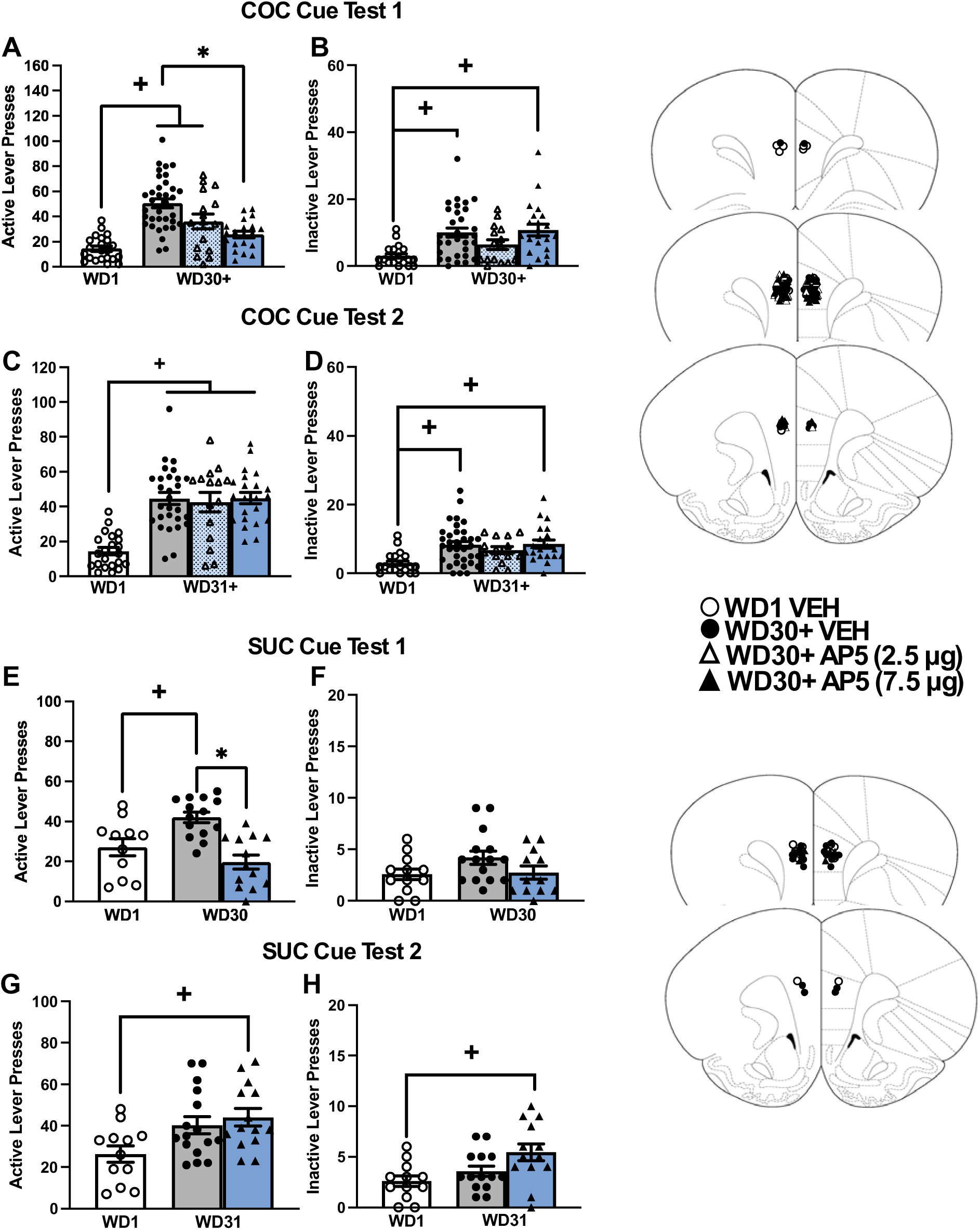
Effects of an intra-PL infusion of D-AP5 on cue-elicited cocaine- and sucrose-seeking during protracted withdrawal. Summary of the effects of microinjection with vehicle (VEH), 2.5 µg or 7.5 µg/side D-AP5 on the number of active **(left)** and inactive **(right)** lever-presses emitted during an initial test for cue-elicited cocaine-craving conducted in later withdrawal (WD30+). **(A)** Relative to early withdrawal controls (WD1-VEH), WD30+-VEH males and WD30+-2.5 AP5 males exhibited incubated craving [for VEH: t(26)=4.435, p<0.001; for 2.5 µg D-AP5: t(36)=3.912, p<0.001], the magnitude of which was reduced by the higher D-AP5 dose [t(53)=4.321, p<0.001], down to the level of WD1 controls [t(29)=1.942, p=0.062]. When carry-over effects were examined the following day (WD31+) (**C**), the attenuating effect of 7.5 µg/side D-AP5 on active lever-pressing was no longer present [t(48)=.046, p=0.964], and all three groups exhibited incubated responding versus WD1-VEH controls [for VEH: t(48)=6.999, p<0.001; for 2.5 µg D-AP5: t(35)=5.355, p<0.001; for 7.5 µg D-AP5: t(42)=7.925, p<0.001]. **(B,D)** Relative to WD1-VEH controls, rats infused with either VEH or 7.5 µg/side D-AP5 exhibited higher inactive lever-pressing on both cue test days than WD1-VEH controls [Test 1, for VEH: t(51)=4.101, p<0.001; for 7.5 µg AP5: t(41)=4.221, p<0.001; Test 2, VEH: t(53)=3.880, p<0.001; 7.5 µg AP5: t(39)=4.126, p<0.001]. Summary of the results from a comparable study of incubated sucrose-seeking, comparing the effects of VEH versus the 7.5µg/side dose of D-AP5 during protracted withdrawal (WD30). **(E)** WD30-VEH controls expressed incubated sucrose-craving [t(27)=4.889, p<0.001], the magnitude of which was reduced by 7.5 µg D-AP5 [t(12)=5.158, p<0.001] to the level of rats tested in early withdrawal [t(22)=1.324, p=0.199], but no group differences were detected for inactive lever-responding during the initial test **(F).** When carry-over effects were examined (WD31), incubated craving was not longer apparent in VEH-infused controls [t(22)=1.324, p=.12], while rats infused with 7.5 µg D-AP5 exhibited higher active **(G)** [t(24)=3.033, p<.001] and inactive **(H)** [t(23)=2.852, p<0.001] lever-presses vs. WD1-VEH controls. Cartoons depicting the placements of the microinjectors within the PL in both experiments are provided. Data represents the means± SEMs of the number of individual rats indicated. *p<0.05 vs. VEH-WD30; +p<0.05 vs. WD1 (withdrawal day 1).

#### NMDAR subunits in the PL

We detected no group differences in the total protein expression of GluN1, GluN2A, and GluN2B subunits within male PL **(Fig.1B-D)** or in GluN1 and GluN2A subunits within female PL **(Fig.1E,F)**. GluN2B tended to be higher on WD30 in female PL, irrespective of cocaine experience **(Fig.1G)** [Withdrawal effect: F(1,47)=1.380, p=0.05], with no significant interaction.

#### CaMKII activity in the PL

CaMKII expression in male PL was lower on WD30, irrespective of cocaine experience **(Fig.1H)** [Withdrawal effect: F(1,42)=21.884, p<0.001]. In contrast, PL CaMKII levels were higher overall in cocaine-experienced females **(Fig.K)** [Group effect: F(1,47)=6.034, p=0.018]. Cocaine-experienced males (**Fig.1I,J**) and females (**Fig.1L,M**) both exhibited higher total and relative p(Thr286)-CaMKII expression within the PL on WD30, corresponding to incubated craving [Group X Withdrawal interactions, total p-CaMKII: for males, F(1,39)=4.177, p=0.049; for females, F(1,44)=4.753, p=0.035; relative p-CaMKII: for males, F(1,40)=7.874, p=0.008; for females, F(1,43)=4.468, p=0.041].

### Intra-PL D-AP5 decreases both incubated cocaine- and sucrose-craving

When the NMDAR competitive antagonist D-AP5 was infused into the PL on WD30, the higher D-AP5 dose (7.5 µg/side) significantly removed the incubation of cocaine-seeking [Group effect: F(1,90)=10.150, p<.001]. This D-AP5 effect was transient as D-AP5 pretreated rats exhibited incubated levels of responding the next day (WD31; **Fig.2C**) [Group effect: F(2,88)=13.922, p<.001]. Neither D-AP5 dose reduced inactive lever-responding on either cue test (**Fig.2B,D**) [Group effect, for WD30: F(3,86)<6.785, p<.001; for WD31: F(3,85)=6.185, p<.001].

The “anti-incubation” effect of intra-PL D-AP5 (7.5 µg/side) generalized from cocaine to sucrose (**Fig.2A**); D-AP5 infusion removed the incubated sucrose-craving (**Fig.2E**) [Group effect: F(2,40)=12.663, p<.001]. In contrast, no D-AP5 effect was detected on inactive lever-pressing on WD30 (**Fig.2F**) [F(2,40)=3.214, p=.052]. When tested the next day, incubated responding had extinguished in WD30-VEH rats, whereas rats infused with D-AP5 now manifested elevated levels of active lever-responding (**Fig.2G**) [Group effect: F(2,42)=4.587, p=.017], as well as high responding on the inactive lever, suggestive of generalized hyperactivity (**Fig.2H**) [F(2,41)=4.603, p=.016]. Thus, intra-PL NMDAR inhibition transiently blocks both incubated cocaine- and sucrose-craving.

### Intra-PL myr-AIP decreases incubated cocaine-craving in a manner independent of NMDARs

To determine the role for CaMKII activation in the expression of incubated craving, distinct groups of rats were infused intra-PL with either the CaMKII inhibitor myr-AIP alone or in combination with 7.5 µg/side D-AP5. Relative to WD1-VEH controls, W30+-VEH rats exhibited incubated cocaine-craving that was blocked by an intra-PL infusion of AP5, myr-AIP, and their combination and the effect of inhibitor co-infusion was greater than either inhibitor alone (**Fig.3A**) [Group effect: F(4,71)=14.05, p<0.0001]. When tested the next day in the absence of any further pretreatment, incubated craving was apparent in WD30+-VEH controls, as well as both D-AP5- and myr-AIP-pretreated rats, but not in rats pretreated with both inhibitors (**Fig.3C**) [Group Effect: F(4,66)=3.913, p=0.0065]. In contrast, no pretreatment affected inactive lever-pressing during the initial Cue Test [Group effect: F(3,51)=2.396, p=0.0790] (**Fig.3B**), whereas inactive lever-responding was lower in myr-AIP pretreated rats, relative to both WD30+-VEH controls and rats with prior D-AP5 pretreatment on the second Cue Test (**Fig.3D**) [Group Effect: F(4,67)=4.261, p=0.004].

**Figure 3:**
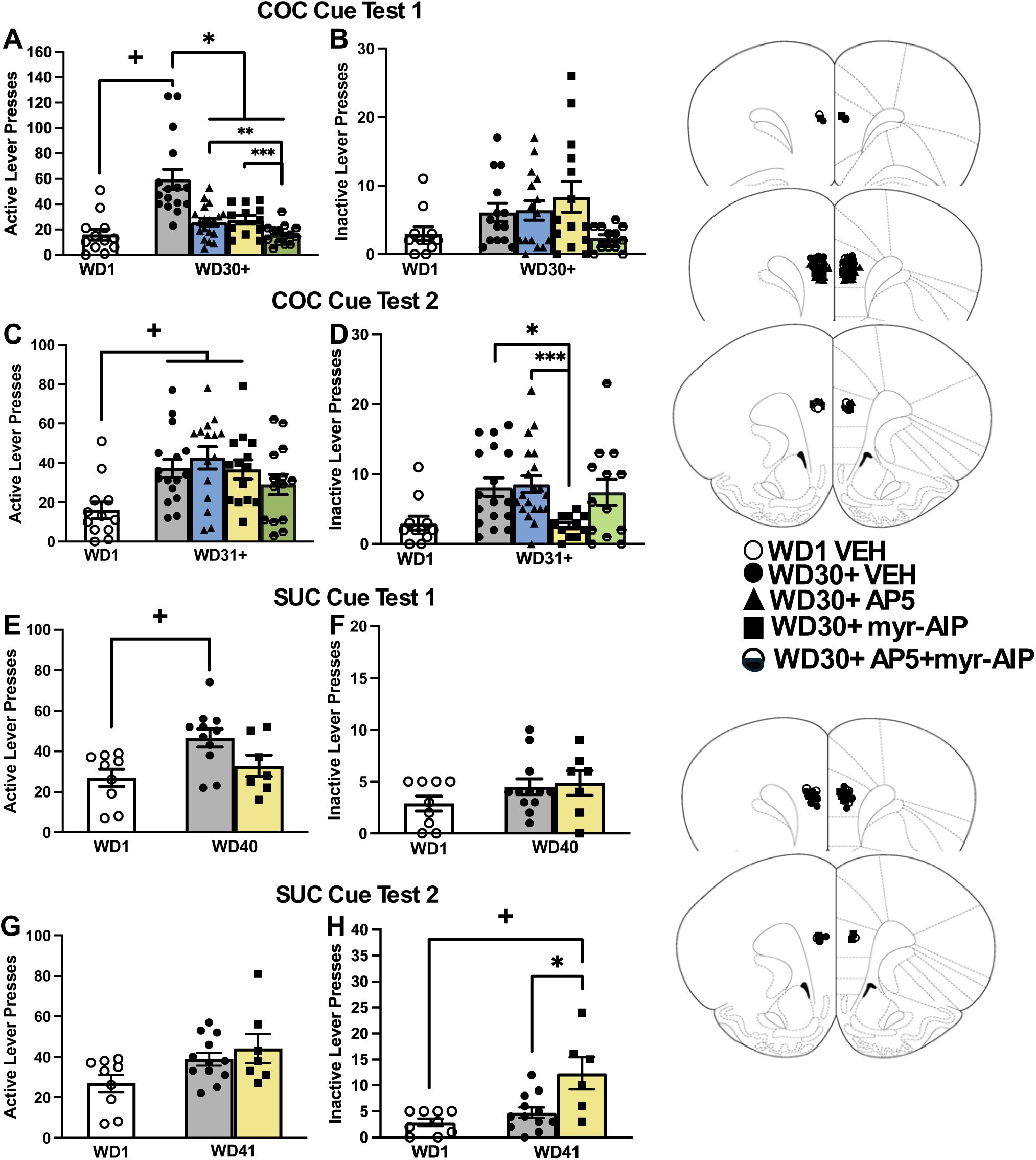
Effects of an intra-PL infusion of D-AP5, myr-AIP, and their combination on cue-elicited cocaine-& sucrose-seeking during protracted withdrawal. Summary of the immediate effects of microinjection with vehicle (VEH), 7.5µg/side D-AP5, 10 pg/side myr-AIP or a combination of the two inhibitors on the number of active **(left)** and inactive **(right)** lever-presses emitted during an initial test for cue-elicited craving conducted in early (WD1) or later withdrawal (WD30+ or WD40), as well as a test for carry-over effects conducted the following day. **(A)** WD30+-VEH rats exhibited incubated cocaine-craving [t(71)=6.189, p<0.0001], that was reduced by intra-PL infusion of D-AP5 [t(71)=5.186, p<0.001], myr-AIP [t(71)=4.298, p=0.0005] and their combination [t(71)=6.447, p<0.0001], with all 3 pretreatments reducing active lever-responding to the level of WD1-VEH controls [for D-AP5: t(29)=1.942, p=0.062; for myr-AIP: t(21)=1.851, p=0.078; for combination: t(24)=0.016, p=0.988]. Inhibitor co-infusion also reduced incubated responding to a greater extent than either AP5 [t(71)=2.907, p=0.0065] or myr-AIP alone [t(71)=3.231, p=0.0034], but did not impact inactive lever-pressing (t-tests, p>0.05) **(B)**. **(C)** When tested the next day, incubated cocaine-craving was apparent in all WD30+ rats, with the exception of the rats with prior co-infusion [WD30+-VEH: t(66)=2.954, p=0.0151; D-AP5: t(66)=3.647, p=.0019; myr-AIP: t(66)=2.794, p=0.0232; co-inhibitor: t(66)=1.759, p=0.2337]. **(D)** Rats pretreated with myr-AIP emitted fewer inactive lever-presses, relative to both WD30+-VEH controls [t(67)=2.918, p=0.0469] and rats with prior D-AP5 pretreatment [t(67)=3.109, p=0.0272]. In a follow-up sucrose study, VEH-infused rats exhibited incubated sucrose-craving on WD40 [t(18)=3.128, p=0.0058], while that myr-AIP-infused rats was not statistically different from either VEH control **(E)** [vs. WD40-VEH: t(16)=1.944, p=0.0697; vs. WD1-VEH t(14)=.8942, p=0.3863]. **(F)** No group differences were apparent for inactive lever pressing on the first Cue Test. When tested the next day, no group differences were detected for active lever-pressing **(G)**, while prior myr-AIP infusion increased inactive lever-pressing relative to both VEH groups **(H)** [vs. WD1-VEH: t(13)=3.566, p=0.0034; vs. WD41-VEH: t(16)=2.948, p=0.0095]. Cartoons depicting the placements of the microinjectors within the PL in both experiments are provided. Data represents the means± SEMs of the number of individual rats indicated. +p<0.05 vs. WD1, *p<0.05 vs. WD30+VEH, ** p<0.05 vs. D-AP5, *** p<0.05 vs. myr-AIP

Intra-PL myr-AIP infusion also blocked the expression of incubated sucrose-craving (**Fig.3E**) [Group effect: F(2,24)=5.211, p=0.0132], without altering inactive lever-responding (**Fig.3F**) [Group effect: F(2,25)=1.3, p=0.2694]. No carry-over effect of myr-AIP infusion was observed for active lever-pressing (**Fig.3G**) [Group effect: F(2,27)=1.8, p=0.1698], while inactive lever-responding was higher in myr-AIP pretreated rats, relative to both VEH groups (**Fig.3H**) [Group effect: F(2,24)=8.9, p=0.0012]. Thus, inhibiting CaMKII activation within the PL is also sufficient to transiently block incubated cocaine- and sucrose-seeking, without overt off-target effects.

### Intra-PL D-AP5 and myr-AIP indiscriminately increase responding in early withdrawal

In stark contrast to their impact on incubated responding, intra-PL infusions of D-AP5 or myr-AIP increased the numbers of active (**Fig.4A,C**) and inactive (**Fig.4B,D**) lever-presses emitted, relative to VEH controls during the early Cue Test [for D-AP5, active lever: t(19)=2.113, p=0.048, inactive lever: t(18)=2.626, p<.001; for myr-AIP, active lever: t(17)=2.813, p=0.0119; inactive lever: t(16)=2.587, p=0.0199]. Thus, the imapcts of of D-AP5 and myr-AIP is selective for the incubated state.

**Figure 4:**
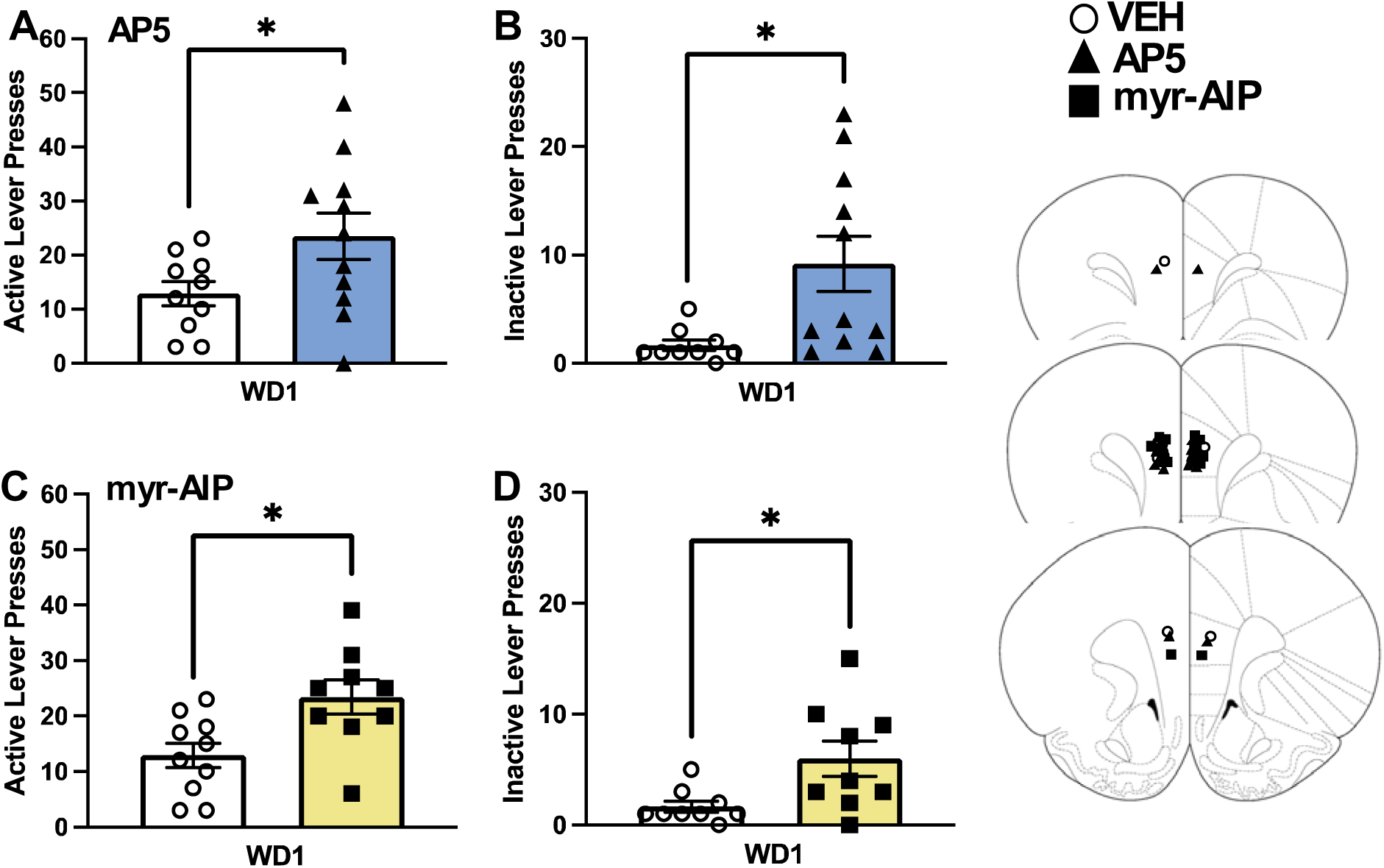
Effects of an intra-PL infusion of D-AP5 or myr-AIP on cue-elicited cocaine-seeking in early withdrawal. Summary of the effects of an intra-PL infusion of vehicle (VEH), D-AP5 (7.5µg/side) **(top)** and myr-AIP (10 pg/side) **(bottom)** on active **(left)** and inactive **(right)** lever pressing on a test for cue-elicited cocaine-seeking conducted on withdrawal day 1 (WD1). Both D-AP5 and myr-AIPincreased both active (**A**) [for D-AP5: t(19)=2.113, p=0.048; for myr-AIP: t(17)=2.813, p=0.0119] and inactive lever-pressing **(B)** [D-AP5: t(16)=2.587, p=0.019; myr-AIP: t(16)=2.587, p=0.0199]. Cartoons depicting the placements of the microinjectors within the PL in both experiments are provided. The data represent the means ± SEMs of the number of individual rats indicated. *p<0.05 vs. VEH.

### Protein correlates of incubated sucrose-craving

As both NMDAR and CaMKII inhibition blocked incubated sucrose-craving **(Fig.2-3)**, we immunoblotted for NMDAR subunit expression and CaMKII activation within the PL and IL of male and female rats tested for cue-elicited craving on WD1 or WD30. The cue-elicited sucrose-craving results are summarized in **Fig.5A** and detailed in [32] with the PL protein expression results provided below and those for IL are in the Supplementary Material and depicted in **Suppl. Fig.3**.

**Figure 5:**
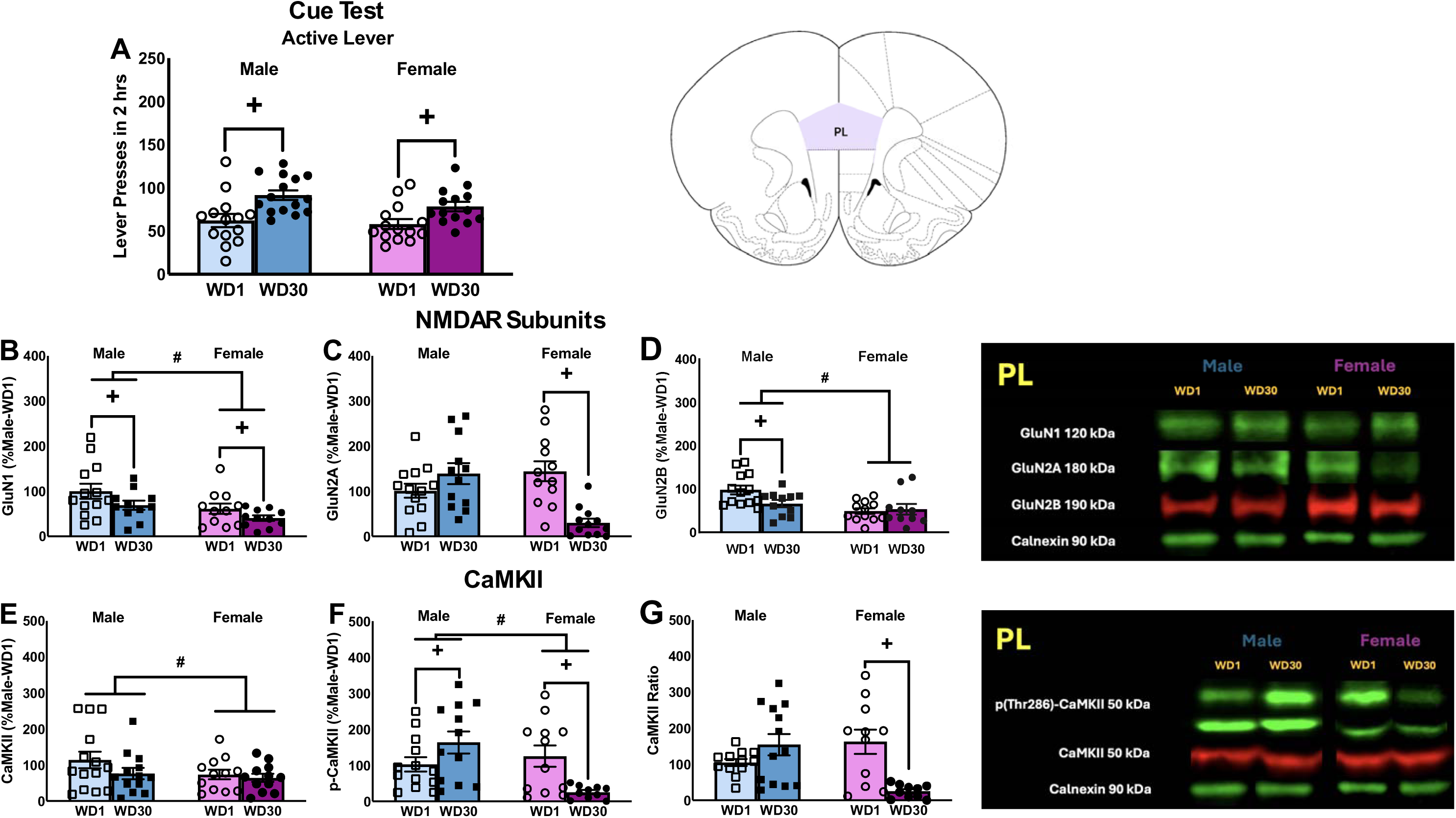
Immunoblotting conducted on the PL subregion of sucrose-incubated rats. **(A)** Active lever-pressing of male and female rats tested for incubated sucrose-craving prior to immunoblotting. The data represent the means ± SEMs of the number of rats indicated and details of the statistical analyses are provided in Ref. 32. Immunoblotting indicated: **(B)** a Sex and Withdrawal effect for GluN1; **(C)** a Sex x Withdrawal interaction for GluN2A that reflected a time-dependent decrease in GluN2A in female [t(22)=4.334, p<0.001], but not male [(22)=1.722, p=0.099], rats**; (D)** a Sex effect and a Sex x Withdrawal interaction for GluN2B, reflecting a time-dependent decrease in GluN2B within the PL of male [t(23)=2.034, p=0.054], but not female rats[t(22)=0.787, p=0.440]; **(E,F)** Sex effects for both CaMKII and p(Thr286)-CaMKII; and **(G)** a Sex x Withdrawal interaction for relative p(Thr286)-CaMKII expression that reflected a time-dependent decrease in female [t(19)=3.792, p=0.001], but not male [t(23)=1.529, p=0.140] rats. Representative immunoblots are provided for all proteins examined. The data represent the means ± SEMs of the individual animals indicated. *p<0.05 vs. Control (main Drug effect); +p<0.05 vs. WD3 (Withdrawal effect).

In contrast to cocaine, we detected a number of group differences in NMDAR expression within the PL of sucrose-experienced rats, some of which coincided with incubated craving. Overall, the expression of GluN1, GluN2B, CaMKII, and p(Thr286)-CaMKII was lower in females versus males (**Fig.5**) [Sex effects, for GluN1: F(1,46)=6.494, p=0.015; for GluN2B: F(1,49)=8.028, p=0.007; for CaMKII: F(1,48)=6.911, p=0.012; for p(Thr286)-CaMKII: F(1,47)=7.967, p=0.007]. Incubated sucrose-craving was associated with increased GluN1 in rats of both sexes (**Fig.5B**) [Withdrawal effect: F(1.46)=4.990, p=0.031], decreased GluN2A in females (**Fig.5C**) [Sex x Withdrawal interaction: F(1,48)= 17.911, p<0.001], decreased GluN2B in males (**Fig.5D**) [Sex x Withdrawal interaction: F(1,49)=4.185, p=0.047] and decreased relative p(Thr286)-CaMKII expression in females (**Fig.5F**) [F(1.46)=14.859, p<0.001]. Thus, the incubation of sucrose-craving is associated with NMDAR subunit expression and CaMKII activation that are both sex-selective and distinct from those observed in cocaine-incubated rats.

## DISCUSSION

Herein, CaMKII hyper-activation within the PL correlates with incubated cocaine-seeking, independent of sex or changes in total NMDAR subunit expression. In contrast, incubated sucrose-seeking molecular correlates exhibit sex-selective changes in both NMDAR subunits and CaMKII activation within both mPFC subregions. Despite these differences, PL NMDAR and CaMKII activity is necessary for incubated craving for both reinforcers and these signaling molecules appear to operate independently within this brain region to drive incubated cocaine-craving.

### NMDAR expression and incubated craving

As reported following short-access IV cocaine self-administration [30], NMDAR subunit expression did not vary in the PL or IL of rats in the present study [Fig.1, Suppl. Fig.1]. Thus, in contrast to dmPFC [18,19], vmPFC NMDAR subunits appear to be less sensitive to gross, withdrawal-induced, changes in whole-cell expression even in rats with considerable cocaine experience. Notably, increased dmPFC GluN2A expression is not observed until 60 days into cocaine withdrawal [18] and similar changes in specific NMDAR subunits within vmPFC may also manifest post-induction of incubation (i.e. after 30 days). Alternatively, incubated cocaine-craving may relate to more subtle shifts in subcellular localization, phosphorylation and/or changes in other NMDAR subunits not examined herein [20,40,41]. The regulation of NMDARs during cocaine withdrawal is complex as exemplified in the only other published study relating NMDARs to incubated cocaine-craving [20]; incubated craving was associated with persistently enhanced NMDAR transmission within the NAc that reflected a switch from GluN2B- to GluN3-containing receptors over the course of protracted withdrawal and the only change in NMDAR subunit expression detected was an increase in the cell surface expression of GluN3A at 48 days into withdrawal [20]. Although both GluN3A and GluN3B are expressed in cortex [42–44], we could not reliably detect these subunits in our tissue and so their relationship to incubated craving is still unknown.

Opposite cocaine [5,45,46], incubated sucrose-craving is associated with reduced AMPA/NMDA ratios within the NAc of adult male rats [47]. Herein, incubated sucrose-craving related to gross changes in PL NMDAR subunit expression [Fig.5], but these changes (lower GluN1 in both sexes; lower GluN2A in females) are predictive of reduced, rather than enhanced, NMDAR transmission in this subregion [15,16]. These same sucrose-incubated females also exhibit lower mGlu5 expression [32] and elevated GluA2/GluA1 ratios within the PL [14], both of which are predicted to blunt calcium-dependent intracellular signaling in this subregion. In contrast, sucrose-incubated males exhibit increased mGlu5 within both vmPFC subregions [32] and elevated IL GluA1 expression [14], possibly indicative of higher CP-AMPAR expression [45]. Taken together, these gross changes in total glutamate receptor expression suggest that incubated sucrose-craving reflects blunted calcium-mediated signaling within the PL of females, but augmented signaling within both mPFC subregions in males. In PFC, all three ionotropic glutamate receptors (iGluRs) are expressed not only on principle glutamatergic projection neurons, but also on GABAergic interneurons, the antagonism of the latter disinhibits pyramidal cell firing [48–50]. As the immunoblotting procedures employed herein cannot discern between different cell types, we do not know if these sex-related changes in glutamate receptor expression are ubiquitous or cell type-specific, to explain how blocking AMPAR/KARs [14] and NMDARs [Fig.2] within the PL all reduce incubated sucrose-seeking to a similar extent in both male and female rats. Nevertheless, the present immunoblotting results for NMDAR subunits provide additional evidence that the biochemical correlates of incubated cocaine- and sucrose-craving by male and female rats are distinguishable in terms of both presynaptic [12] and postsynaptic [14,18,30,32,33] aspects of glutamate transmission.

### CaMKII activation and incubated craving

Replicating and extending the results from our earlier study [30], CaMKII activation was increased selectively within the PL of cocaine-incubated male and female rats [Fig.1], while cocaine-incubated males also exhibited lower CaMKII activation within the IL [Suppl. Fig.1]. In both cases, CaMKII activation was independent of gross changes in either AMPAR [14] or NMDAR subunit expression [Fig.1; Suppl. Fig.1]. The present results for the IL differ slightly from our earlier study of incubated cocaine-craving [30], in which IL CaMKII activation was either lower or unchanged at both withdrawal time-points in cocaine-experienced males. Although CaMKII activation varied with estrous cycle phase coincident with incubated responding [Suppl. Fig.2], overall, cocaine-experienced females exhibited no overt changes in IL CaMKII activation despite exhibiting robust incubated cocaine-craving [Suppl. Fig.1]. Taken all together, we posit that incubated cocaine-craving correlates more reliably with higher CaMKII activation within the PL, rather than within the IL. As CaMKII activation within neither subregion aligned with AMPAR or NMDAR expression, changes in CaMKII activation are likely driven by either more subtle adaptations that impact receptor function and/or perhaps other upstream affectors [28,48]. Given our neuropharmacological results to date implicating the activation of all three major iGluRs within the PL in incubated cocaine-craving, an important goal of future work is to identify more precisely which iGluR subtypes are stimulating CaMKII to drive the cocaine-incubated state.

As observed for NMDAR subunits, incubated sucrose-craving was associated with sex-dependent changes in indices of CaMKII activation within both the PL and the IL [Fig.5; Suppl. Fig.3]. In the PL, males exhibited higher total p(Thr286)-CaMKII expression, while this phospho-kinase was lower in females [Fig.5]. The lower PL CaMKII activation of sucrose-incubated females coincides with lower GluN1 and GluN2A expression [Fig.5], implicating blunted NMDAR-mediated CaMKII activation within the PL in incubated sucrose-craving by female rats. As such a mechanism is at odds with our neuropharmacological results [Fig.2,3], we posit that the time-dependent down-regulation of NMDAR-CaMKII signaling within the PL of sucrose-incubated females may reflect adaptations within GABAergic interneurons and a disinhibition of corticofugal glutamatergic projections. The relationship between vmPFC CaMKII activation and NMDAR expression in males is less straight-forward. Sucrose-incubated males exhibit a “cocaine-like” increase in CaMKII activation within the PL that is inversely related to GluN1 expression [Fig.5]. These males also exhibit higher CaMKII activation within the IL, unrelated to changes in total NMDAR [Suppl. Fig.3] or AMPAR [14] subunits. However, increased CaMKII activation within the IL does coincide with increased indices of mTOR and PKCε activation, as well as mGlu5 expression as reported previously in these same males [32], implicating calcium-dependent signaling through mGlu5 as potentially upstream of CaMKII activation within the IL of males. While the IL is purported to inhibit cocaine-seeking in the extinction-reinstatement model of relapse [51,52], there is growing appreciation that its function in reinforcer-seeking is more nuanced [53,54]. Our present results, coupled with the complex regulation of cocaine-seeking by manipulations of endogenous glutamate within the IL [13], implicate neuroadaptations within the IL as potentially contributing to incubated craving, particularly for palatable food reinforcers.

### NMDAR and CaMKII inhibition within the PL blocks incubated craving

Cocaine cue-elicited glutamate release within the PL is necessary for the expression of incubated cocaine-craving [12,13], while sucrose cue-elicited glutamate release declines during protracted sucrose abstinence [12]. Despite this and distinctions in the protein profiles within vmPFC associated with incubated cocaine-versus sucrose-craving [14,30,32; present study], we recently identified PL AMPAR/KARs as important for the expression of incubated craving for both reinforcers [14] and the present results for D-AP5 [Fig.2] extend this regulation to NMDARs. Further, we identify CaMKII as a potential intracellular mediator of both incubated cocaine-craving and sucrose-craving [Fig.3]. A direct role for NMDAR-CaMKII signaling within the PL in driving incubated craving is supported, in part, by evidence that inhibitor co-infusion also blocked incubated cocaine-craving [Fig.3]. However, the magnitude of the “anti-incubation” effect of inhibitor co-infusion was larger than either inhibitor alone [Fig.3], arguing that other upstream CaMKII activators are also involved. Group1 mGlu receptors are not likely candidates as intra-vmPFC antagonist infusions do not affect the expression of incubated craving [35]. However, CP-AMPARs or -KARs could provide a source of intracellular calcium that might act synergistically with NMDARs to activate CaMKII. Indeed, CP-AMPARs are upregulated within vmPFC of cocaine-sensitized mice [52], a large body of work implicates NAc CP-AMPARs in incubated cocaine-craving [5,20,45,46] and AMPAR/KAR activation within PL is necessary for cocaine- and sucrose-craving [14]. As such, a goal of future work is to assess the role of intra-PL CP-AMPAR/KAR subtypes both alone and in combination with myr-AIP on incubated cocaine- and sucrose-craving.

As observed for the AMPAR/KAR antagonist NBQX [14], intra-PL infusion of either D-AP5 or myr-AIP induced elevated indiscriminate lever-pressing in rats during early withdrawal [Fig.4]. It is currently unclear if these effects reflect the induction of perseverative lever-pressing [55], stereotyped behavior [56,57] or learning/memory impairments [21–23]. However, the fact that intra-PL iGluR inhibition induces opposing effects on cue-elicited responding between early and later withdrawal argues that (1) the “anti-incubation” effects of intra-PL infusion of iGluR and CaMKII inhibitors do not likely reflect ataxia or off-target motivational deficits and (2) time-dependent changes occur within iGluR and/or CaMKII signaling pathways within PL during cocaine withdrawal that alter their behavioral pharmacology.

In contrast to the protracted “anti-incubation” effects observed following an intra-vmPFC infusion of PI3K [11] and PKCε inhibitors [10], the effects of intra-PL infusion of AMPAR/KAR [14], NMDAR [Fig.2] and CaMKII [Fig.3] inhibitors on incubated cocaine- and sucrose-craving are transient; in both cases, cue-elicited responding returned to the levels of incubated VEH-infused controls the day following microinjection. Thus, although iGluRs and CaMKII are highly implicated in learning and memory [21–24], their inhibition within the PL do not appear to facilitate aspects of extinction learning or memory to reduce subsequent cue-reactivity. This said, a modest carry-over effect of D-AP5 and myr-AIP co-infusion was detected the day following treatment t [Fig.3]. Thus, by definition [4], inhibitor co-infusion blocked incubated cocaine-craving for at least 24 h. As CaMKII, PI3K/Akt/mTOR and PKCε signaling interact in complex ways to affect cell function upon iGluR activation [e.g., 58-61], it will be important in future studies to identify the kinase-kinase interactions involved in the neuroplasticity associated with incubated craving as combined pharmacological treatments may have greater efficacy in curbing incubated craving.

### Conclusions and Caveats

Despite no overt sex differences in behavior, incubated cocaine- and sucrose-craving are associated with distinct, often sex-dependent, profiles of GluN1, GluN2A and GluN2B expression and CaMKII within mPFC subregions. Although more work is required to fully appreciate the nuances (e.g. subcellular location, cell type specificity or further subunit analyses) of NMDAR expression and CaMKII activation within mPFC subregions in incubated craving, our neuropharmacological results clearly demonstrate a necessary role for both NMDAR and CaMKII within the PL in incubated cocaine- and sucrose-craving. However, it remains to be determined whether CaMKII or specific NMDAR subtypes operate within IL to regulate incubated craving, particularly in light of the sex-specific changes in IL protein expression observed herein and the apparent relationship between estrous cycle phase, CaMKII activation and the magnitude of incubated craving in cocaine-experienced female subjects. Lastly, the larger and more enduring effect of intra-PL NMDAR and CaMKII co-inhibition implicate at least one other affector of CaMKII in driving incubated craving that may facilitate extinction learning. In sum, while the results of the present study identify NMDARs and CaMKII within the PL as important for incubated cocaine- and sucrose-craving, a considerable amount of work remains to uncover the precise mechanisms through which these signaling molecules operate to augment cue-reactivity during protracted abstinence of relevance to pharmacotherapeutic strategies for treating intensified craving for both drug and food reinforcers.

## Funding and Disclosure

This work was supported by NIH/NIDA grant R01DA053328 (KKS) and R01DA51100 (TEK). Additional research support was provided through the University of California Santa Barbara’s Faculty Research Assistance Program (KKS) and Undergraduate Creative and Research Activities program (NMS, SRC, MGT, RMK, HHTD, and TLL). LLHS is supported, in part, by a NSF-AGEP CA HSI Alliance Fellowship and the NIH BRAIN and Blueprint DSPAN Award F99NS141388. FJC is supported, in part, by a Bill and Melinda Gates Predoctoral Scholarship. The authors have nothing to disclose.

## Author Contributions

The authors contributed to this report in the following ways. Conceptualization, LLHS, FJC, NMS, SRC, MGT, RMK, HHTD, TLL, TEK and KKS; formal analysis, KKS and LLHS; investigation, LLHS, NMS, SRC, JEB, AYN, MGT, MLM, RMK, CRL, HHTD, ABL, TCC, FJC; writing—original draft preparation, LLHS and KKS; writing—review and editing, LLHS, NMS, SRC, JEB, AYN, MGT, MLM, RMK, CRL, HHTD, ABL, TCC, FJC, TEK and KKS; visualization, LLHS and KKS; supervision, TEK and KKS; project administration, LLHS and KKS; funding acquisition, KKS, TEK, LLHS, FJC, NMS, SRC, MGT, RMK, HHTD and TLL. All authors have read and agreed to the published version of the manuscript.

## Notes

### Competing Interest Statement

The authors have declared no competing interest.

