## Supplemental Material for "Inactivation of NMDAR and CaMKII signaling within the prelimbic cortex blocks incubated cocaine- and sucrose-craving"

**SUPPLEMENTAL MATERIALS**

**Methods**

*Subjects.* Adult male and female Sprague Dawley rats (225-250 g; Charles River Laboratories, Hollister, CA, USA), were housed under a reverse light-dark cycle (lights off:10:00 am), with *ad libitum* access to food and water. All procedures aligned with the guidelines of the NIH Guide and Care and Use of Laboratory Animals (NIH publication No. 80-23, revised 2014), with approval from the University of California Santa Barbara Institutional Animal Care and Use Committee under protocol 829. The rats employed for immunoblotting were the same as those described in recent reports for incubated cocaine- [14] and sucrose-craving [32] and thus, no new rats were required to obtain the immunoblotting data described herein.

*Surgery.* Rats employed in the neuropharmacological studies of incubated cocaine-craving that were slated for testing in early withdrawal (WD1) underwent both IC and IV surgical procedures on the same day during a single surgical session, while rats slated for testing in late withdrawal (WD30+) underwent IV catheterization prior to cocaine self-administration procedures and IC surgery 7 days prior to their WD30+ date to minimize subject attrition due to cannulae occlusion. Rats employed in the immunoblotting study of incubated cocaine-craving underwent IV surgical procedures only.  All surgical procedures were performed under isoflurane anesthesia (4% induction, 1-3% maintenance; Covetrus, Portland, ME). For IV catheterization, a chronic polyurethane catheter (12 cm long; 0.023 inner diameter, 0.038 in outer diameter; Instech Laboratories, Plymouth Meeting, PA) was inserted into the right jugular vein and tunneled subcutaneously along the shoulder to a back inclusion where it was connected to a threaded, protruding 22-gauge metal guide cannula (P1 Technologies, Roanoke, VA), within a rat infusion harness (Instech Laboratories, Plymouth Meeting, PA). The cannula was capped with both a plastic and a metal cap to prevent infection and immediately flushed with 0.1 ml of sterile cefazolin (100 mg/ml) and 0.1 ml of sterile heparin (70 U/ml).  Bilateral cannulae were implanted above the PL (AP: +3.0; ML: ±0.75, DV: -2.00 mm from Bregma) and secured to the skull with dental acrylic. Note that rats slated for testing in early withdrawal underwent the bilateral cannulation surgery immediately following IV catheterization and 7 days of post-operative care.  To minimize subject attrition due to clogged guide cannulae, rats slated for testing in later withdrawal underwent intracranial cannulation 7 days prior to cue testing, followed by 4 days of post-operative care.  For 48 hours post-procedure, rats were injected subcutaneously with Meloxicam (2 mg/kg) once a day to alleviate pain and inflammation. For rats slated for cocaine self-administration, post-operative care also consisted of daily IV infusions of cefazolin and heparin to maintain catheter patency. Catheter patency was confirmed just before cocaine self-administration training by administration of 0.1 ml of sodium Brevital (10 mg/ml) IV by loss of muscle tone.

*Cocaine Self-administration and Sucrose Reinforcement.* Rats were trained to self-administer IV cocaine (5-second IV infusion of 0.25mg/kg/0.1ml; MilliporeSigma, Burlington, MA) or 45 mg banana-flavored sucrose pellets (BioServ, Flemington, NJ) for 6hr/day over 10 days. In both cases, an active lever-press resulted in reinforcer delivery and a 20-second tone-light stimulus complex (78 dB, 2 kHz) signaling that delivery. Any active lever presses that occurred during the 20-second stimulus delivery were recorded, but led to no subsequent infusion or pellet delivery. To prevent potential overdose in cocaine trials, rats were limited to a maximum of 100 infusions for the first day. After rats completed 10 days of self-administration, they were slated to undergo testing on either 1 day (WD1) or following at least 30 days of withdrawal (WD30+). During the periods of withdrawal, rats remained housed in the colony room and continued to receive *ad libitum* access to food and water. Any rats emitting less than ten active lever presses in the last three days of self-administration training were dropped from the study (did not undergo withdrawal or testing) and not included in the final statistical analyses of the results.

*Immunoblotting.* PL and IL tissue was homogenized in a lysis buffer solution consisting of 10 mL RIPA Buffer (Boston Bioproducts, Milford, MA), 0.2 M sodium orthovanadate, 0.144 M sodium fluoride, 100 µL  Phosphatase Inhibitor Cocktail 3 (MilliporeSigma, Burlington, MA), and a complete Mini-Tab Protease Inhibitor Cocktail Tablet (Roche Diagnostics, Mannheim, Germany). Homogenates were then centrifuged at 10,000 RPM for 20 min and the supernatant of the homogenates were stored at 80°C. Protein samples (19 𝜇l/lane) were subjected to SDS-PAGE on Tris-acetate gradient gels (3–8%) (Invitrogen), followed by wet polyvinylidene difluoride (MilliporeSigma, Burlington, MA) membrane transfer and membranes were preblocked with TBS containing 0.1% (v/v) Tween 20 and 5% (w/v) nonfat dried milk powder or BSA for a minimum of 1 h before overnight incubation with primary antibody. The following rabbit antibodies were used: GluN1 (1:250 dilution; Cell Signaling Technology; 5704S), GluN2A (1:250 dilution; MilliporeSigma; 07-632), p(Thr286)-CaMKII (1:1000 dilution; Cell Signaling Technology; 3361); GluN3A (1:150 dilution; Antibodies Incorporated; N416/40); GluN3A (1:150 dilution; MilliporeSigma; 07-356). The following mouse antibodies were also used: GluN2B (1:1000 dilution; Invitrogen; MA1-2014) and  CaMKII (1:1000 dilution; Millipore; 05-532). Calnexin was used to control for protein loading and transfer (1:1000 dilution; Enzo Life Sciences; ADI-SPA-860). Membranes were washed with TBST, rinsed with TBS, and incubated in either a goat anti-rabbit IRDye 800CW secondary antibody (1:10,000 dilution; Li-Cor; 925-3221) or a goat anti-mouse IRDye 680RD secondary antibody (1:10,000 dilution; Li-Cor; 925-68070) and imaged in an Odyssey Fc Infrared Imaging System (Li-Cor Biosciences, Lincoln, NE, USA). Protein expression was then quantified using Image Studio, raw values were normalized to the corresponding Calnexin signal, and then averaged to their control groups (WD3-Control for the cocaine study; WD1-Male for the sucrose study). As neither GluN3A antibody reliably detected this subunit in our tissue, the data for GluN3A not presented.

*Statistical Analyses*.  Consistent with our recent study of AMPA receptor correlates of incubated cocaine-craving [14], the data were normalized to the average of the two or three cocaine-naive Control-WD3 animals on each membrane and analyzed using a Group (Control vs. Cocaine) X Withdrawal (WD3 vs. WD30) ANOVA, separately for male and female subjects. The immunoblotting study of incubated sucrose-craving did not include a sucrose-naive control [32]. Thus, the samples from males and females were immunoblotting concurrently, the data normalized to the average of the three Male WD1 rats on each gel and then analyzed using a Sex X Withdrawal (WD1 vs. WD30) ANOVA. To relate protein expression to estrous phase, the immunoblotting data were analyzed using a Phase (estrus, diestrus, proestrus; no rats were found to be in metestrus) X Withdrawal ANOVA [14]. For the neuropharmacological studies of incubated craving, the average number of active and inactive lever presses emitted during either Cue Test was analyzed using a Treatment (WD1-VEH, WD30+-VEH, WD30+-AP5, WD30+-myr-AIP, WD30+-AP5+myr-AIP) X Sex ANOVA. For the neuropharmacological experiment conducted on WD1, the  data were analyzed using a Treatment (VEH vs. drug)  X Sex ANOVA, separately for D-AP5 and myr-AIP as these drugs were examined in distinct experiments. Significant main effects or interactions were further investigated with t-tests or tests for simple effects. As the results of the statistical analyses failed to indicate any main Sex effects or interactions (p>.999), the data were collapsed across male and female subjects in all experiments. Outliers were identified and excluded from the analyses using the ± 1 × IQR rule; however, in instances where too many outliers were identified, we adopted the ± 3 × IQR rule to ensure that only the most extreme outliers were removed. Alpha was set to 0.05. IBM SPSS Statistics software (version 29.0 for Macintosh) was used for all statistical tests, and GraphPad Prism software (version 9.3.1 for Macintosh) was used to create all graphs.

**Results**

**NMDAR subunit expression and CaMKII activation in the IL of rats exhibiting an incubation of cocaine-craving.** Similar to the PL, analyses of NMDAR subunits in the IL failed to detect any changes in NMDAR subunits in male rats  **(Suppl. Fig.1A-C)** [for GluN1: F(1,44)<0.026, p>0.873; for GluN2A: F(1,44)<0.020, p>0.888; for GluN2B: F(1,44)<0.002, p>0.968], and no differences were observed for GluN1 and GluN2B subunits in the IL of females **(Suppl. Fig.1D,E)** [for GluN1: F(1,44)<2.447, p>0.126; for GluN2B: F(1,47)<0.134, p>0.716]. A Withdrawal effect was detected for the GluN2A subunit [F(1,47)=10.386, p=0.002] within the IL of female rats, but this effect did not vary as a function of cocaine history as indicated by no significant interaction **(Suppl. Fig.1F)**.

An examination of total CaMKII expression within the IL detected a Group effect [F(1,45)=5.748, p=0.021] in male **(Suppl. Fig.1G),** but not female, rats [all F(1,43)<0.115, p>0.736] **(Suppl. Fig.1J),** which reflected higher protein expression in cocaine-experienced males. No group differences were detected for p(Thr286)-CaMKII within this subregion in either males **(Suppl. Fig.1H)** or females [for males: F(1,45)<1.497, p>0.228; for females:  F(1,44)<1.582, p>0.216] **(Suppl. Fig.1K)**. In male rats, we detected a Group effect in relative p(Thr286)-CaMKII expression [F(1,45)=8.748, p=0.005], which reflects lower expression in cocaine-experienced males **(Suppl. Fig.1I)**. In female rats, we detected no changes in relative CaMKII within the IL [F(1,43)<0.115, p>0.736] **(Suppl. Fig.1L)**

**Influence of estrous phase on the expression of NMDAR subunits and CaMKII in incubated cocaine-craving.** Previous research demonstrates that the magnitude of incubated cocaine-craving can be modulated by estrous phase [65-67] and within these same rats, we have shown that estrus females demonstrate higher responding on the active and inactive lever, than those in other phases of the estrous cycle [14; see **Suppl. Fig.2A**]. Within the present study, we found that this estrus effect on cocaine-seeking behavior was not associated with changes in GluN1, GluN2A, or GluN2B expression within the PL (**Suppl. Fig.2B-D**) [GluN1: F(1,27)<0.387; p>0.684, GluN2A:  F(1,27)<0.293, p>0.749; GluN2B: F(1,27)<0.384 p>0.685]. Similarly, we detected no changes associated with estrous phase in the expression of GluN1, GluN2A, or GluN2B  within the IL (**Suppl. Fig.2H-J**) [GluN1: F(1,27)<0.646; p>0.534, GluN2A:  F(1,27)<0.164, p>0.690; GluN2B: F(1,27)<0.242, p>0.787]. Within the PL, there was no effect of estrous phase on CaMKII, p(Thr286)-CaMKII, or relative CaMKII (**Suppl. Fig.2E-G**) [CaMKII: F(1,27)<0.184; p>0.834, p(Thr286)-CaMKII:  F(1,27)<0.122, p>0.730; CaMKII ratio: F(1,27)<0.007, p>0.993]. Finally, within the IL, we detected no effect of estrous phase on CaMKII (**Suppl. Fig.2K-L**) [F(1,27)<0.007, p>0.993] or p(Thr286)-CaMKII [F(1,27)<0.168, p>0.846]. There was a significant Withdrawal x Stage effect on relative p(Thr286)-CaMKII expression (**Suppl. Fig.2M**) [F(1,27)<5.504, p=0.012] and deconstruction along the Stage factor revealed a time-dependent increase in the relative p(Thr286)-CaMKII in estrus females [estrus: t(5)=2.525, p=0.053; diestrus: t(13)=1.602, p=0.133; proestrus: t(4)=1.416, p=0.230].

**NMDAR subunit expression and CaMKII activation in the IL of rats exhibiting an incubation of sucrose-craving.** Females exhibited higher expression of GluN1(F(1,55)=7.490, p=0.008] in the IL (**Suppl. Fig.3A**), but no group differences were observed for GluN2A [F(1,55)<2.209, p>0.205] or GluN2B subunits [F(1,55)<1.557, p>0.218] (**Suppl. Fig3B-C**). Further, there were no changes in the expression of CaMKII within the IL **(Suppl. Fig.3D)** [F(1,56)<0.092, p>0.763]. We detected a significant Sex effect [F(1,56)=9.955, p=0.003], Withdrawal effect [F(1,56)=4.691, p=0.035], and Sex x Withdrawal interaction [F(1,56)=9.027, p=0.004] for p(Thr286)-CaMKII, the latter of which reflected a time-dependent increase in p(Thr286)-CaMKII in male, but not female, rats **(Suppl. Fig.3E)** [for Males:  t(26)=2.924, p=0.007, for Females: t(26)=0.897, p=0.378]. Similarly, a Sex effect [F(1,54)=13.198, p<0.001], Withdrawal Effect [F(1,54)=11.335, p=0.001], and Sex x Withdrawal interaction [F(1,54)=19.688, p<0.001] were detected for relative p(Thr286)-CaMKII expression that reflected a time-dependent increase selectively in males [for Males: t(24)=4.652, p=0<0.001, for Females:  t(26)=0.941, p=0.355]. Representative immunoblots are provided for all proteins examined. The data represent the means ± SEMs of the individual animals indicated. *p<0.05 vs. Control (main Drug effect);  +p<0.05 WD1 vs. WD30 (incubation).

**Supplemental Figure Legends**

**Supplemental Figure 1:  Immunoblotting in the IL subregion of cocaine-incubated rats.** Summary of the changes in NMDAR expression and CaMKII activation within the IL of cocaine-naive (Control) or -experienced (Coc) male and female rats. No group differences were detected in NMDAR subunit expression within the IL of male rats **(A-C)**, while a Withdrawal effect was detected for GluN2A in the IL of females **(D-F)**. Cocaine-experienced males exhibited lower relative p(Thr286)-CaMKII expression, but no other group differences were detected for CaMKII-related measures in males **(G-I)** or for any CaMKII-related measure within the IL of females **(J-L).** Representative immunoblots are provided for all proteins examined. The data represent the means ± SEMs of the individual animals indicated. *p<0.05 vs. Control (Cocaine effect); +p<0.05 WD3 vs. WD30 (withdrawal effect).

**Supplemental Figure 2:  Immunoblotting in mPFC subregions of cocaine-incubated female rats across the estrous cycle.** Summary of how the number of active and inactive lever-presses varied as a function of estrous cycle **(A)** (P=proestrus; E=estrus; D=diestrus; no females were found to be in metestrus). No relationship was observed between estrous cycle phase and the expression of NMDAR subunits within the PL **(B-D)** or CaMKII activation in this subregion **(E-G)**. No estrous cycle effects were detected for the IL expression of NMDAR subunits **(H-J)**, although estrus females exhibited a time-dependent increase in the relative expression of p(Thr286)-CaMKII **(I-K)**. The data represent the means ± SEMs of the individual animals indicated. +p<0.05 WD3 vs. WD30 (incubation).

**Supplemental Figure 3:  Immunoblotting in the IL subregion of sucrose-incubated rats.** Females exhibited higher GluN1 expression within the IL **(A)**, but no group differences were observed for GluN2A **(B)**, GluN2B **(C)**, or CaMKII **(D)**. For both the total **(E)** and relative **(F)** expression of p(Thr286)-CaMKII within the IL, only males exhibited a time-dependent increase in phospho-kinase expression. Representative immunoblots are provided for all proteins examined. The data represent the means ± SEMs of the individual animals indicated. *p<0.05 vs. Males (Sex effect);  +p<0.05 WD1 vs. WD30 (incubation).

**Suppl. Figure 1**

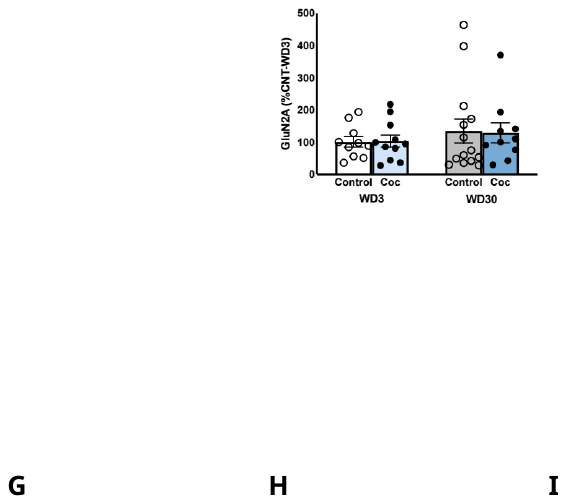

**Suppl. Figure 2**

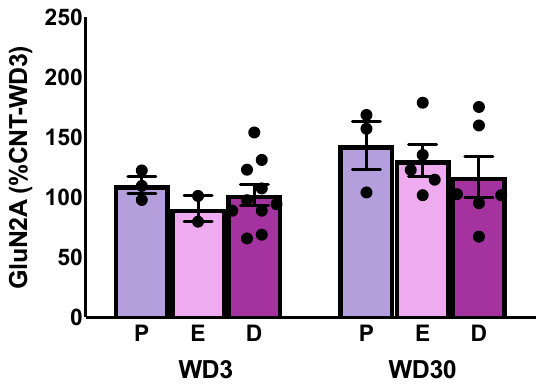

**Suppl. Figure 3**

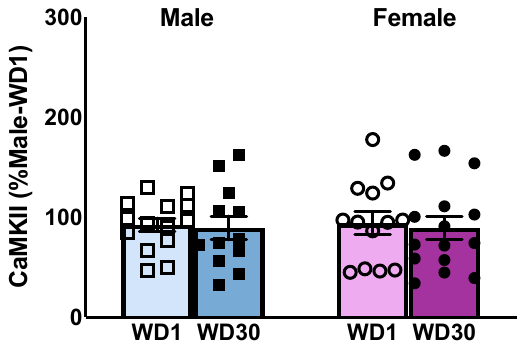
